# An Effective Metal Nanoparticle-Based Drug Delivery System for an *In Vitro* Model of Non-Small Cell Lung Cancer

**DOI:** 10.64898/2026.06.05.730380

**Authors:** Natalia Piergies, Magdalena Oćwieja, Katarzyna Pogoda, Agnieszka Panek, Maciej Roman, Kamil Raszka, Wojciech M. Kwiatek

**Affiliations:** Institute of Nuclear Physics Polish Academy of Sciences, PL-31342 Krakow, Poland; Jerzy Haber Institute of Catalysis and Surface Chemistry Polish Academy of Sciences, Niezapominajek 8, PL-30239 Krakow, Poland; SOLARIS National Synchrotron Radiation Centre, Jagiellonian University, Czerwone Maki 98, 30-392 Krakow, Poland

## Abstract

This study presents the development and spectroscopic characterization of an erlotinib-functionalized gold nanoparticle (erlotinib:AuNP) nanosystem designed for targeted delivery to metastatic non-small cell lung cancer H1299 cells. Initial MTS assays demonstrated that free erlotinib induced a concentration-dependent reduction in cell viability, while 0.1 µM erlotinib exhibited negligible cytotoxicity and was therefore selected for nanosystem fabrication. AuNPs alone showed minimal toxicity toward H1299 cells over the investigated concentration range. Following conjugation of erlotinib with AuNPs, the resulting nanosystems reduced cell viability to approximately 60%, indicating enhanced biological activity of the drug after nanoparticle-assisted delivery. Fluorescence microscopy confirmed the intracellular internalization of the nanosystems in H1299 cells, with nanoparticle aggregates predominantly localized in the perinuclear and perimitochondrial regions. Three-dimensional Raman spectroscopy (3D RS) mapping further verified the intracellular localization of the conjugates through characteristic Raman signatures of erlotinib:AuNPs. Importantly, 3D RS enabled detection of nanosystems at concentrations below the sensitivity limit of fluorescence imaging, demonstrating superior analytical performance for intracellular nanosystem tracking. Atomic force microscopy–infrared (AFM–IR) spectroscopy coupled with principal component analysis (PCA) demonstrated substantial biochemical modifications induced by the erlotinib:AuNP nanosystems, including enhanced lipid-related spectral features and significant alterations in protein secondary structure, particularly the increased contribution of unordered and antiparallel *β*-turn conformations. The obtained results demonstrate that combining plasmonic nanocarriers with advanced vibrational spectroscopy enables highly sensitive monitoring of intracellular drug delivery and nanosystem-induced biochemical responses in cancer cells.

## INTRODUCTION

The concept of the drug delivery system assumes the designing and development of carriers that can effectively increase the local concentration of the drug inside the target cells while reducing the possible penetration of physiological cells.^1,2^ This can limit the bothersome side effects and decrease the costs of the applied therapeutic regimen. The main challenge in this field is to design a drug/carrier system that establishes a stable connection between the drug and the vehicle while exhibiting low or no toxicity, efficient uptake, and controlled release within the cell. The most attractive structures used as carriers include: liposomes, membrane vesicles, and metal nanoparticles.^3–5^ Among them, the metal nanoparticles may not be so prominent, however, these substrates exhibit unique properties that ensure their successful application in many biomedical fields. The main benefit is the unchallenging modification of their surface which allows for appropriate functionalization and therefore more controlled biocompatibility.^6^ Other interesting attributes are associated with the optical properties which ensures possibilities to achieve enormous enhancement of the spectral signal employed in such techniques as: surface–enhanced Raman spectroscopy (SERS) and surface – enhanced infrared absorption spectroscopy (SEIRA).^7,8^ This amplification of the spectral signal is associated with a local electric field induced near the metal surface. This field is strongly anisotropic,^9^ specifically, the component perpendicular to the metal surface is significantly larger than the component tangent to the metal surface. This phenomenon has certain consequences, specifically, vibrations in which the tensor polarizability varies perpendicular to the metal surface, exhibiting the greatest enhancement in SERS spectra.^10,11^

At the same time, the vibrations for which the dipole moment is normal to the surface dominate the SEIRA spectra.^12–14^ Based on the well-known surface selection rules dedicated to the SERS and SEIRA effects, respectively, it is possible to characterize the spatial orientation of the particular functional groups of a drug after its immobilization on the metal nanocarrier.

This is crucial since the conformation of the drug often determines the biological activity. Moreover, the conformation of the drug can be monitored under different conditions such as pH of the environment, temperature, and time elapsed since adsorption. Such an approach also ensures information about the stability of the created drug/nanocarrier connection.^15^

Our previously published data allowed us to perform detailed studies of how erlotinib and gefitinib, the drugs approved in non-small cell lung cancer (NSCLC) target therapy, adsorb on the silver, gold, and platinum nanoparticles, (AgNPs, AuNPs, and PtNPs, respectively).^16–21^ The considerations based on the SERS, and SEIRA combined with the atomic force microscopy (AFM–IR) confirm that the type of the applied metal surface strongly affects the drug adsorption. Namely, erlotinib interacts with the AgNPs mainly through the phenylacetylene ring, forming a very stable connection. This connection is not disturbed by the temperature and even time elapsed since adsorption. For AuNPs the drug/carrier interaction occurs *via* phenylacetylene ring and quinazoline moiety, while for PtNPs this connection is principally due to the quinazoline moiety. In the case of gefitinib, we observe a strong interaction with gold and silver nanoparticles mainly through quinazoline and morpholine moieties.

In this study, we investigated the effects of a drug delivery system based on AuNPs functionalized with erlotinib on H1299 cells, a cell line derived from lymph node metastasis of non-small cell lung cancer (NSCLC). Both biological assays and spectroscopic techniques were employed to evaluate the cellular response to the developed nanosystems. In addition, the cellular uptake and intracellular localization of the AuNP-based drug carriers were assessed in H1299 cells to determine their interaction with cancer cells and their potential as an effective drug delivery platform.

## EXPERIMENTAL

### Cell culture

Human metastatic non-small cell lung cancer cells (H1299) were cultured in RPMI-1640 medium supplemented with 10% fetal bovine serum (FBS). Cells were maintained at 37°C in a humidified atmosphere containing 95% of air and 5% of CO_2_.The medium was refreshed according to the cell growth requirements.

### MTS assay

Cells were seeded into 24-well plates at a density of 1.5 × 10^4^ cells per well and cultured for 24 h under standard conditions. Subsequently, the culture medium was replaced with serum-free DPBS supplemented with Ca^2+^ and Mg^2+^ for 2 h. Following serum starvation, cells were treated with selected concentrations of erlotinib, AuNPs, and erlotinib:AuNP nanosystems. Cell viability was assessed after 72 h of incubation using the MTS assay. Briefly, CellTiter 96® AQueous One Solution reagent was added directly to the wells and incubated for 1 h. The absorbance of the generated formazan product was measured at 490 nm using a Spark 10 M (Tecan) multimode microplate reader. The obtained values were normalized against untreated control cells. All experiments were conducted in triplicate.

### Fluorescent staining

A series of fluorescence staining experiments were performed to evaluate the influence of the applied treatment conditions on the morphology of H1299 cell line. Particular emphasis was placed on assessing alterations in nuclear morphology, mitochondrial organization, and the actin cytoskeleton. Cells were seeded onto circular glass coverslips (Ø = 15 mm, thickness 0.13–0.16 mm) at a density of 4 × 10^4^ cells/mL and cultured for 24 h under standard conditions. Subsequently, the cells were treated with AuNPs at concentrations of 10 mg/L, 1 mg/L, and 0.1 mg/L, as well as with erlotinib:AuNP nanosystems at a concentration of 0.1µM:10 mg/L, 0.1 µM:1mg/L, 0.1 µM:0.1 mg/L. Following 24 h of incubation, mitochondria were stained using 100 nM MitoTracker for 15 min. The cells were then fixed in 4% paraformaldehyde (PFA) solution in PBS for 20 min at room temperature. The actin cytoskeleton was visualized using Alexa Fluor Phalloidin diluted 1:200 in PBS. After three washing steps with PBS, nuclei were counterstained with Hoechst solution (1:10,000). Fluorescence images were acquired using a Leica DMi8 fluorescence microscope equipped with a 100× oil immersion objective. Image processing, visualization, and analysis were performed using ImageJ software.

### 3D Raman spectroscopy mapping

Raman measurements were performed using inVia Raman spectrometer (Renishaw, UK) coupled to a confocal microscope equipped with a 100× objective and a CCD detector. A 633 nm laser was used as the excitation source, with the laser power at the sample surface maintained at approximately 1 mW. 3D Raman spectral maps were acquired for three individual cells per experimental condition, with a spectral resolution of approximately 1 cm^−1^ over the range of 3100–100 cm^−1^. The acquisition time was 1 s per spectrum, and a single accumulation provided a satisfactory signal-to-noise ratio. For each map, five z-stacks were collected with a 500 nm step size in the z-direction and a 1 µm step size in the x- and y-directions. The spectrometer was calibrated using the Raman band of an internal silicon standard. After each measurement, the sample surface was inspected to verify the absence of laser-induced damage.

### AFM–IR spectroscopy measurements

Measurements were carried out using NanoIR2 system (Anasys/Bruker/ Germany) working in the contact mode using silicon gold-coated PR-EX-nIR2 probes by selecting a contact resonance frequency centered at 180 kHz using a Gaussian filter with a half-width of 50 kHz. Infrared excitation was generated using a multichip tunable quantum cascade laser (QCL; MIRcat-QT, Daylight Solutions) operating in the spectral range of 1800–1200 cm^−1^ with a spectral resolution of 2 cm^−1^. For each experimental condition, spectra were acquired from 10 individual cells, with 10 measurement points collected diagonally across each cell. Each spectrum represented the average of 256 excitation pulses. Additionally, AFM topography images were recorded for every analyzed cell using a cantilever scan rate of 0.5 Hz and a spatial resolution of 500 × 500 pixels.

## RESULTS AND DISCUSSION

### The development strategy of the erlotinib:AuNP conjugates as a drug delivery system

Figure 1A illustrates the first step of the applied approach to the preparation of the erlotinib:AuNP system with the ability to effectively transport the drug to cells. The cells were exposed separately to variuos concentrations of erlotinib and AuNPs to identify a non-cytotoxic concentration of erlotinib and to evaluate the potential cytotoxic effects of AuNPs on the studied cell lines. As expected, the MTS assay results demonstrate that with increasing erlotinib concentrations, a reduction in cell viability is observed (Figure 1B). A statistically significant decrease in cell viability by 50% relative to the control was observed at an erlotinib concentration of 50 µM, while an erlotinib concentration of 0.1 µM showed cell viability comparable to the control. Therefore, the latter drug concentration (marked by a red star in Figure 1 B) was used for the nanosystem development. In the case of the AuNPs, the MTS studies reveal a non-significant influence on cell viability. Namely, the higher concentrations of AuNPs (5 mg/L and 10 mg/L) reduce cell viability to ∼90%. In turn, after the treatment with lower AuNPs concentrations, the viability of H1299 cells drops to ∼80 %. This seemingly counterintuitive effect may be attributed to the aggregation of nanoparticles upon their addition to the cell culture medium, which can affect their cellular uptake. Such aggregation is more likely to occur at higher nanoparticle concentrations, potentially reducing the fraction of particles available for internalization and thereby attenuating their cytotoxic effects. This phenomenon could explain the greater reduction in cell viability observed at lower AuNP concentrations.

**Fig. 1.**
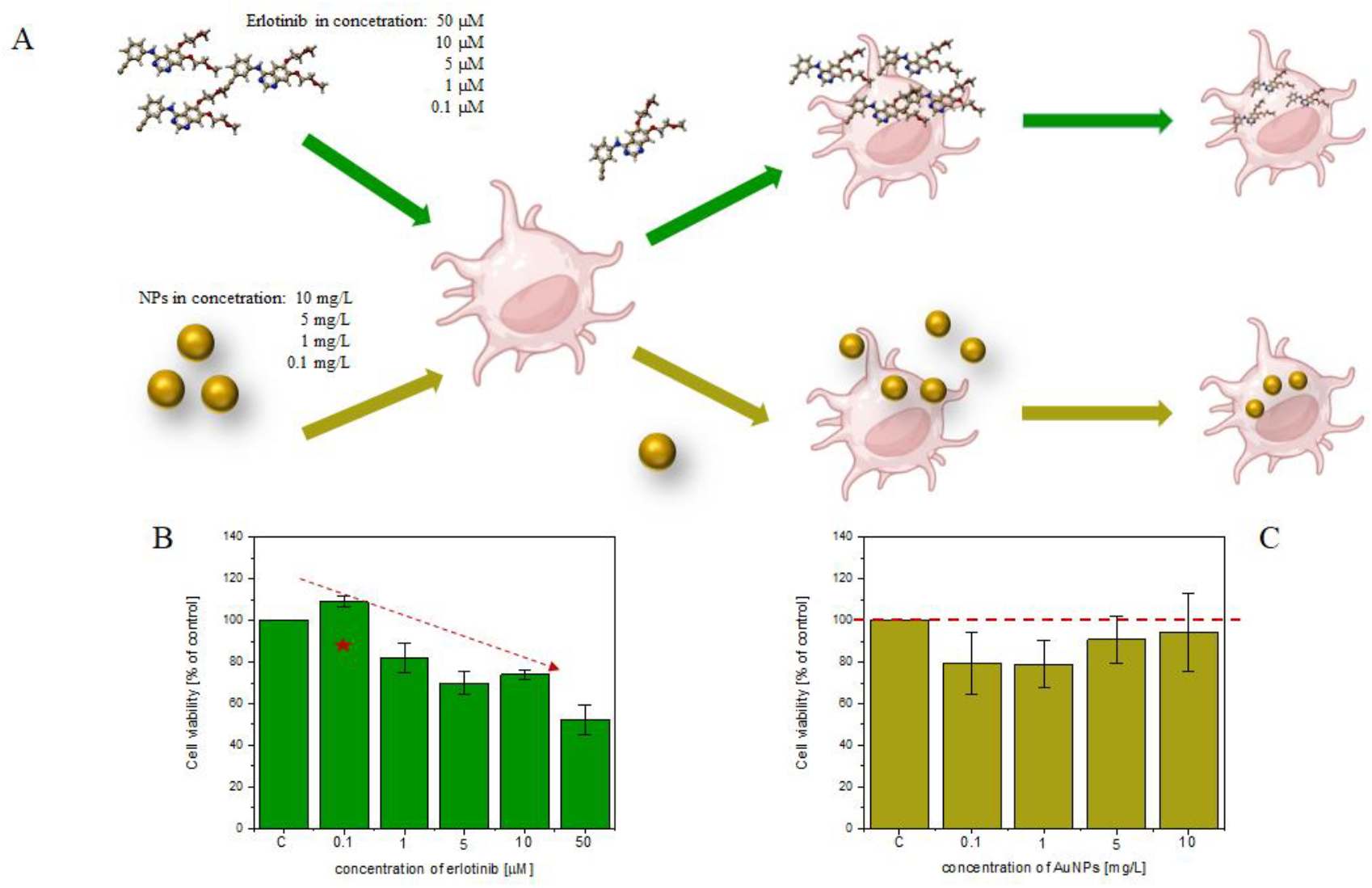
Scheme of experimental procedure of the first step of the conjugates designing (A) together with MTS results of dose-dependent effect of erlotinib (B) and AuNPs (C) on H1299 cell line. The viability of cell lines was determined after 72 h of the treatment by different concentrations of erlotinib (0.1 µM; 1 µM; 5 µM; 10 µM; 50 µM) and AuNPs (0.1 mg/L; 1 mg/L; 5 mg/L; 10 mg/L;), respectively. Cell viability values were normalized to untreated control cells (c). Values represent mean ± SD of nine replicates. The red star indicates the selected erlotinib concentration for the further conjugates development.

Based on the results described above, erlotinib and AuNPs at selected concentrations were used for the preparation of the drug–carrier system. Figure 2A presents the second step of the procedure, in which erlotinib at a concentration of 0.1 µM was deposited on AuNPs at different concentrations, and such prepared nanosystems were used for the cell treatment.

**Fig. 2.**
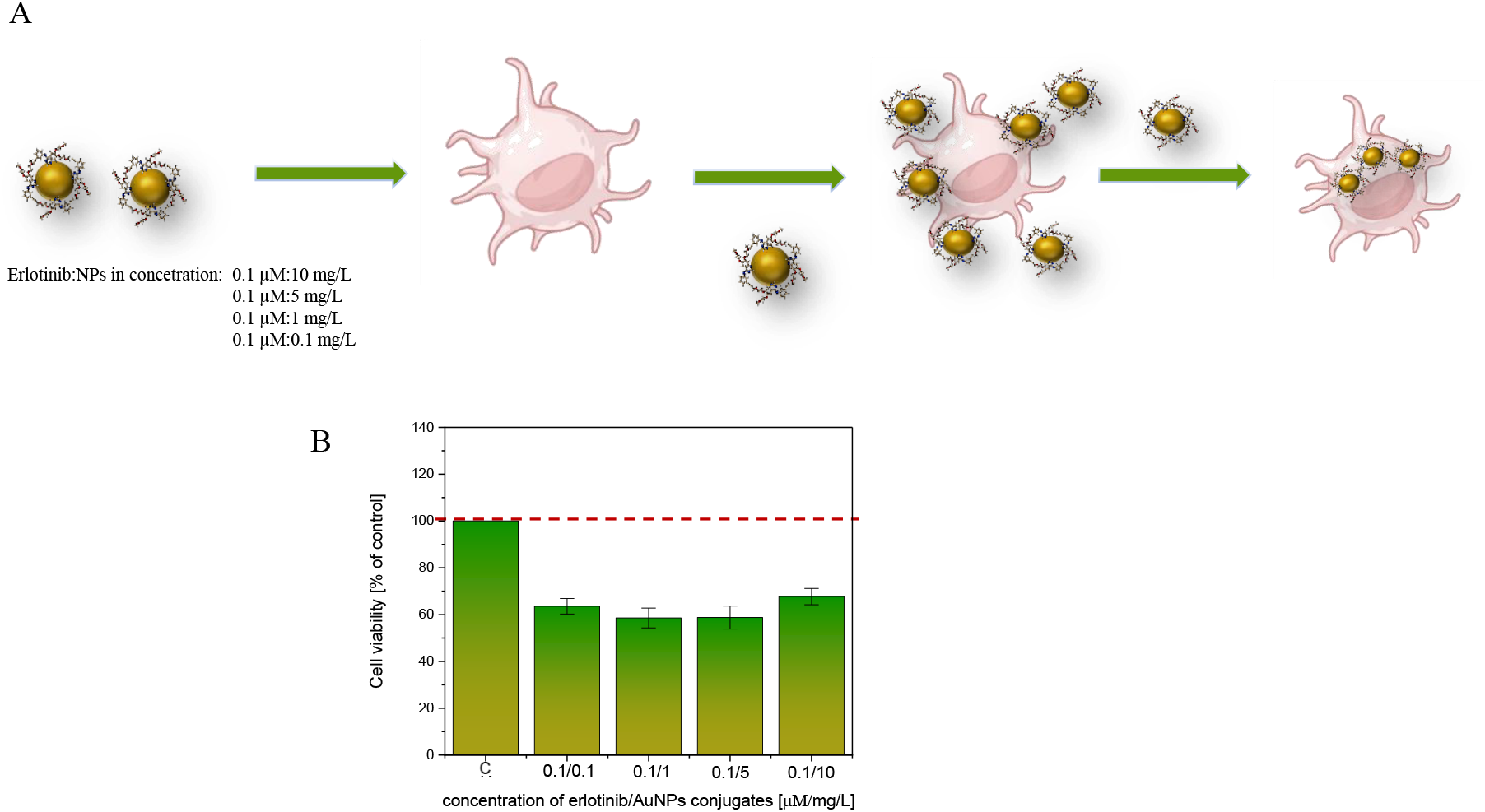
Scheme of experimental procedure of the second step of the conjugates designing (A) together with MTS results of dose-dependent effect of erlotinib:AuNP conjugates (B) on H1299 cell line. The viability of cell lines was determined after 72 h of the treatment by different concentrations of erlotinib:AuNP conjugates (0.1 µM/ 0.1 mg/L; 0.1 µM/1 mg/L; 0.1 µM/5 mg/L; 0.1 µM/10 mg/L), respectively. Cell viability values were normalized to untreated control cells (c). Values represent mean ± SD of nine replicates.

According to the MTS data, the applied treatment reduces the cell viability to ∼60% (Fig. 2B). Importantly, no significant changes in cell viability were observed as the concentration of AuNPs in the nanosystems increased. This suggests that the main toxic effect comes from the drug activity rather than from the influence of the AuNPs.

### Fluorescent imaging

To confirm the cellular internalization of the developed erlotinib:AuNP nanosystems, fluorescence staining was performed. Figure 3 presents representative fluorescence images of H1299 cells treated with AuNPs at a concentration of 10 mg/L, 1 mg/L, and 0.1 mg/L as well as with erlotinib:AuNP systems at concentrations of 0.1 µM:10 mg/L, 0.1 µM:1 mg/L, and 0.1 µM:0.1 mg/L, respectively. Following treatment, the cells were stained with fluorescent probes to visualize actin filaments, mitochondria, and cell nuclei. The blue fluorescence corresponds to nuclei, green fluorescence indicates actin, and red fluorescence marks mitochondria. Dark aggregates observed in the fluorescence images of cells exposed to AuNPs and erlotinib:AuNP systems represent the distribution of AuNPs, confirming successful intracellular uptake of the nanosystems. Notably, these structures appear to localize at the perinuclear and perimitochondrial regions. Such aggregates were not detected in control cells that were not exposed to nanoparticles or conjugates. Furthermore, no dark intracellular aggregates characteristic of AuNP accumulation were observed in cells treated with AuNPs at concentrations of 1 mg L^−1^ and 0.1 mg L^−1^ (Fig. 3).

**Fig. 3.**
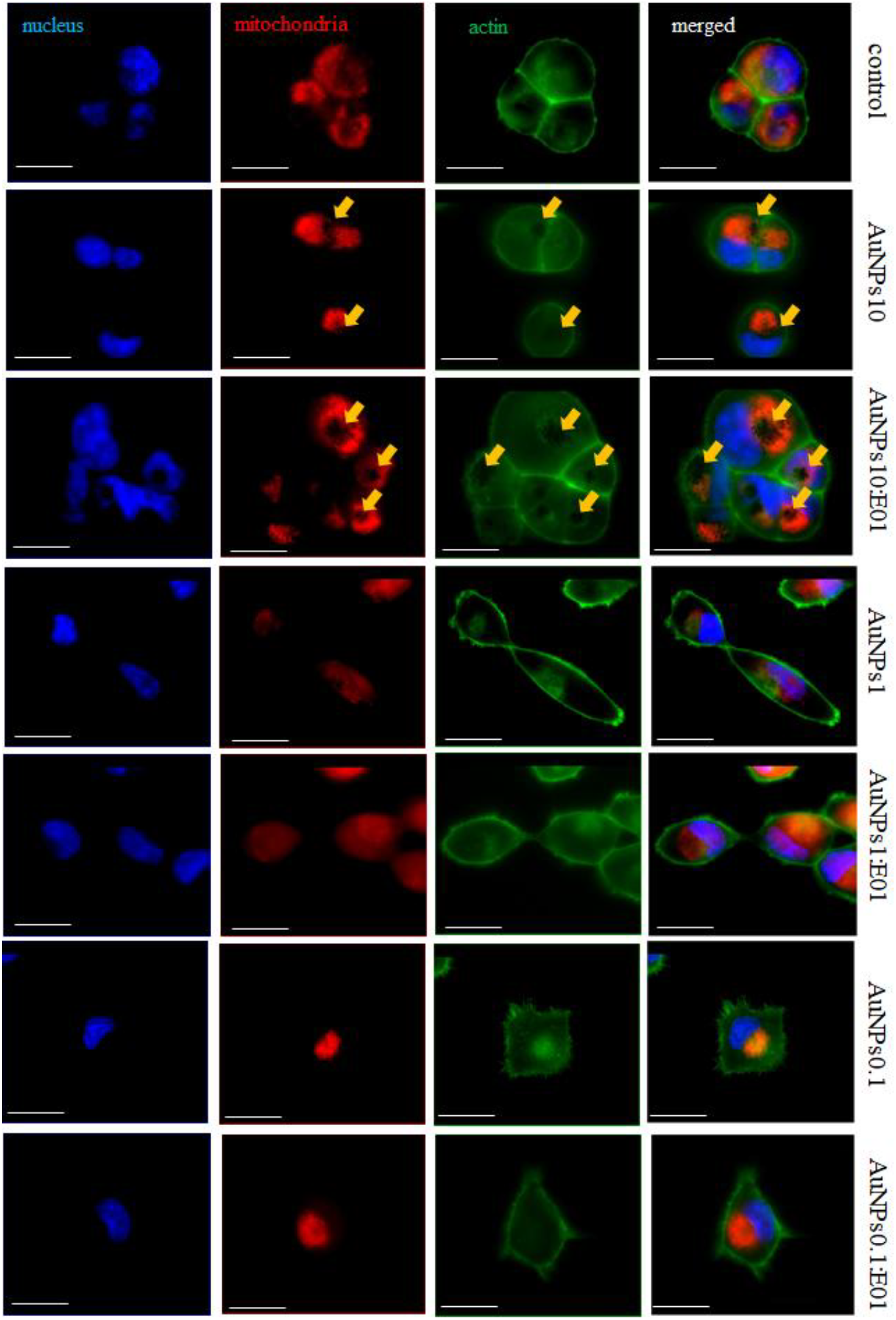
The fluorescent images of H1299 cells untreated (control) and treated with AuNPs and erlotinib:AuNPs conjugates containing 10 mg/L, 1 mg/L, and 0.1 mg/L of AuNPs (AuNPs10, AuNPs1, and AuNPs0.1), and 10mg/L of AuNPs and 0.1 µM of erlotinib (AuNPs10:E01), 1mg/L of AuNPs and 0.1 µM of erlotinib (AuNPs1:E01), 0.1mg/L of AuNPs and 0.1 µM of erlotinib (AuNPs01:E01), respectively, for 24h. The cells were stained – nucleus (blue), mitochondria (red), and actin (green). The arrows indicate the conjugates’ accumulations. Scale bar: 20 µm.

### Spectroscopic studies

To provide additional confirmation of nanosystem internalization, three-dimensional Raman spectroscopy (3D RS) mapping was performed. Since the nanosystem erlotinib:AuNPs exhibit characteristic Raman bands, mainly at approximately 270 cm^−1^ and 2000 cm^−1^,^18^ respectively, cluster analysis was applied to the acquired 3D RS maps. Figures 4 and 5 present a comparison of fluorescence microscopy images and 3D RS maps of H1299 cells treated with erlotinib:AuNP conjugates at concentrations of 0.1 μM erlotinib:10 mg/L AuNPs and 0.1 μM erlotinib:0.1 mg/L AuNPs, respectively. For the conjugates containing 10 mg/L AuNPs and 0.1 µM erlotinib, fluorescence microscopy revealed a distinct intracellular distribution pattern. These observations were further corroborated by 3D RS mapping, which confirmed cellular internalization of the nanosystems and indicated their predominant localization within the perinuclear and perimitochondrial regions. In contrast, conjugates containing 1 mg/L AuNPs and 0.1 μM erlotinib were successfully detected by 3D Raman mapping (Fig. S1; Supporting Information), whereas these structures were not observed in the corresponding fluorescence microscopy images. For nanosystems containing 0.1 mg/L AuNPs and 0.1 μM erlotinib (Fig. 5), no intracellular dark aggregates were detected, however, 3D Raman mapping still enabled the identification and localization of the conjugates within the cells. These results demonstrate the superior sensitivity of Raman imaging for detecting low concentrations of erlotinib:AuNP conjugates in cellular environments. Furthermore, the averaged spectra associated with the clusters identified in the 3D RS maps provided additional evidence for the intracellular localization of the erlotinib-functionalized nanosystems in studied experimental conditions. Figures 4G, 5G, and S1G compare the averaged cluster spectra obtained from the cells (black solid line) with the SERS spectrum of erlotinib adsorbed onto AuNPs at 37 °C (red dotted line). In all cases, characteristic Raman bands of erlotinib were clearly observed in the spectra acquired from the cells by 3D RS mapping. The bands located at approximately 2000 cm^−1^, 1593 cm^−1^, and 994 cm^−1^ were assigned to C≡C stretching, aromatic C=C stretching, and aromatic ring deformation vibrations of erlotinib, respectively.^18^ These results clearly demonstrate that 3D RS mapping provides a significantly lower detection limit compared with fluorescence staining and enables precise tracking of drug-loaded nanosystems within cells.

**Fig. 4.**
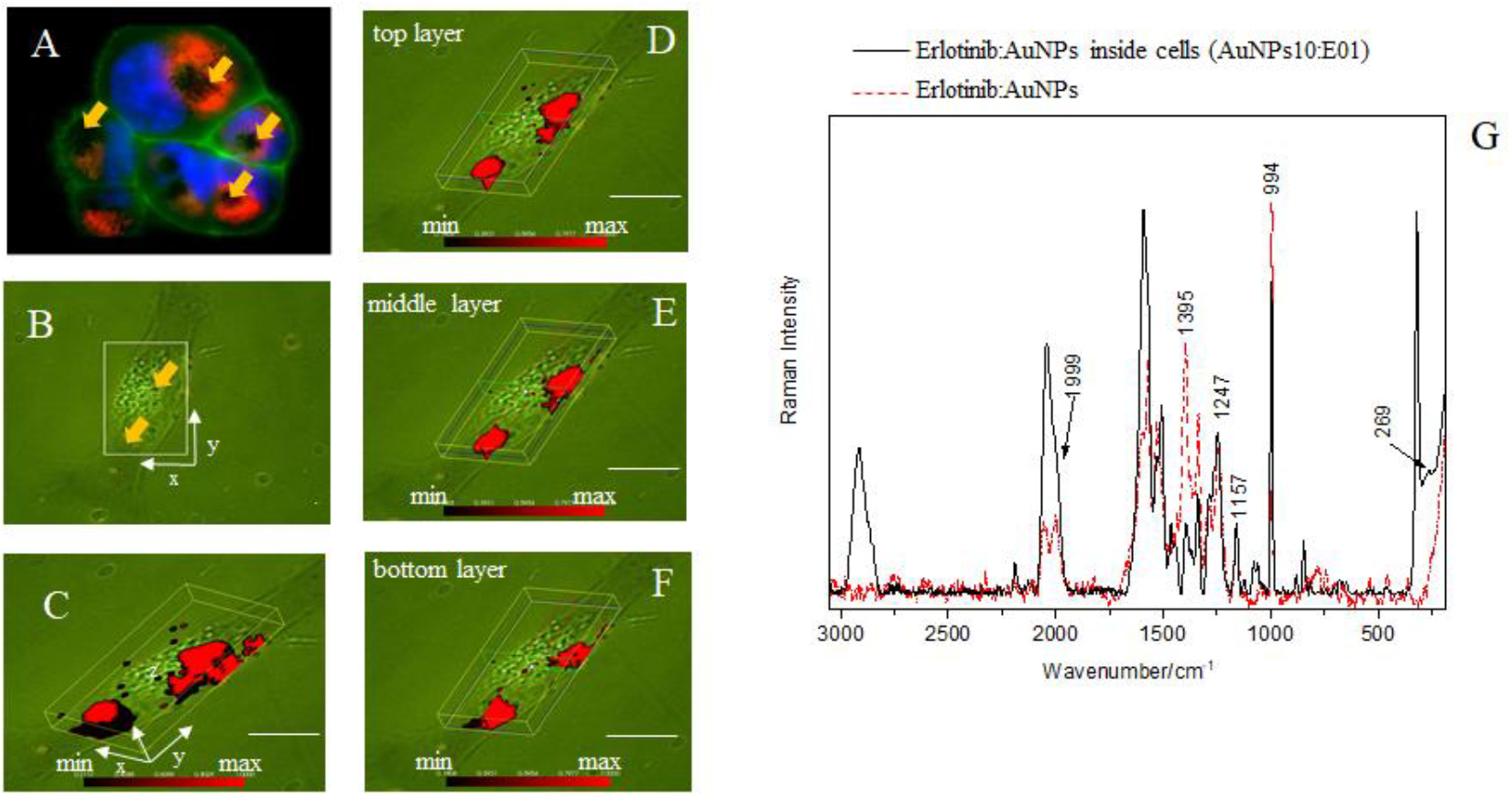
The fluorescent (A) and microscope image (B) of H1299 cells treated for 24 h with AuNP:erlotinib conjugates containing 10 mg/L AuNPs and 0.1 µM erlotinib (AuNPs10:E01). The intracellular distribution of the conjugates, determined by cluster analysis (CA), is shown in panels C–F. Panel G presents the averaged Raman spectrum of the intracellular AuNP:erlotinib cluster (black solid line) compared with the SERS spectrum of the AuNP:erlotinib conjugate recorded after 24 h of adsorption at 37 °C (red dotted line). The arrows indicate the conjugates’ accumulations. Scale bar: 20 µm.

**Fig. 5.**
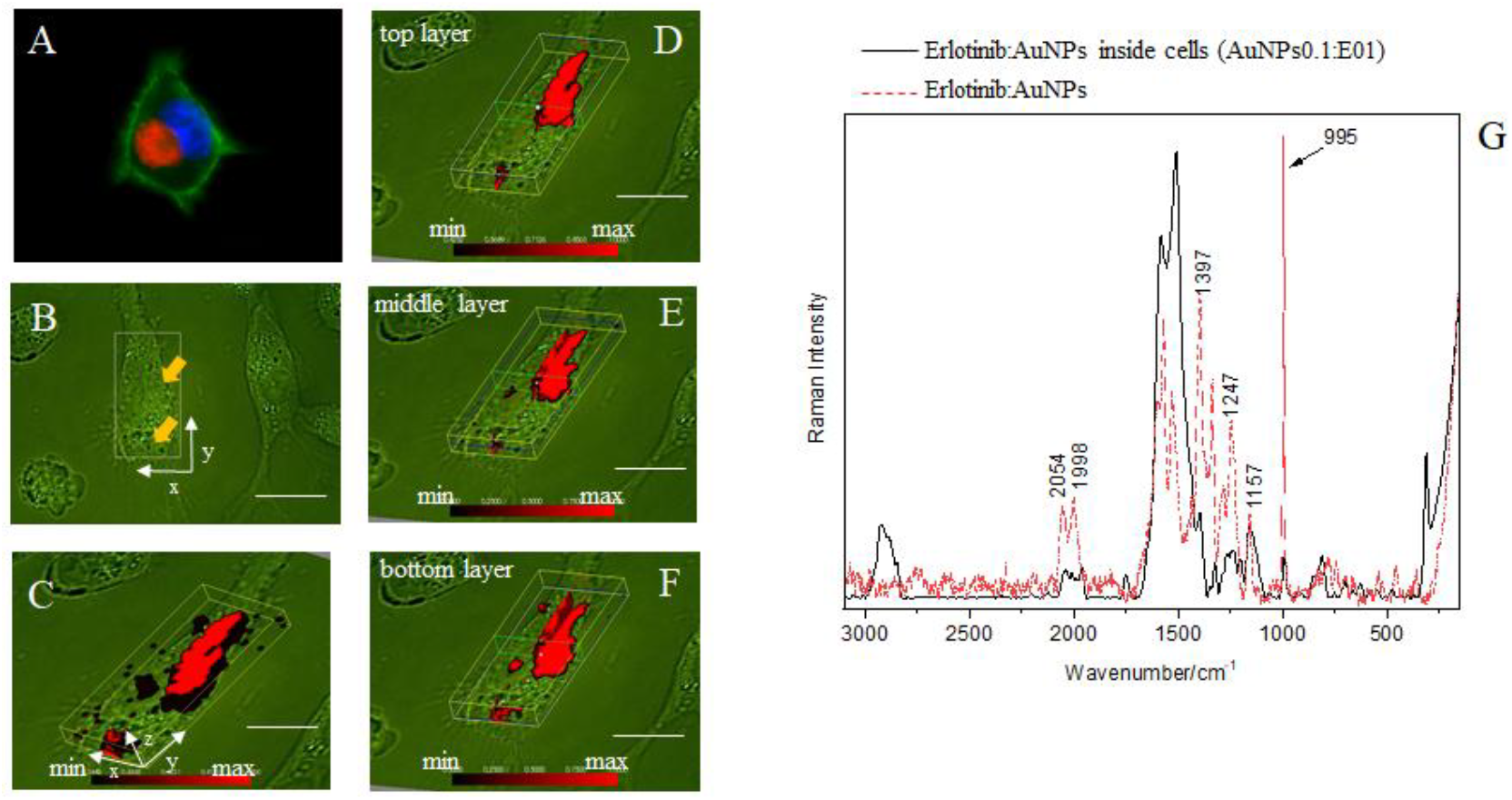
The fluorescent (A) and microscope image (B) of H1299 cells treated for 24 h with AuNP:erlotinib conjugates containing 0.1 mg/L AuNPs and 0.1 µM erlotinib (AuNPs0.1:E01). The intracellular distribution of the conjugates, determined by cluster analysis (CA), is shown in panels C–F. Panel G presents the averaged Raman spectrum of the intracellular AuNP:erlotinib cluster (black solid line) compared with the SERS spectrum of the AuNP:erlotinib conjugate recorded after 24 h of adsorption at 37 °C (red dotted line). The arrows indicate the conjugates’ accumulations. Scale bar: 20 µm.

To investigate the biochemical changes induced by the internalization of the studied nanosystems, AFM–IR measurements were performed. Figure 6 presents representative AFM topographies of H1299 cells together with the measurement profile indicating the locations from which AFM–IR spectra were acquired. The scan size of each AFM image was individually adjusted to match the dimensions of the analyzed cell. Subsequently, principal component analysis (PCA) was applied to the collected AFM–IR spectra to evaluate treatment-induced biochemical alterations. The PCA results for H1299 cells treated with erlotinib, AuNPs, and

**Fig 6.**
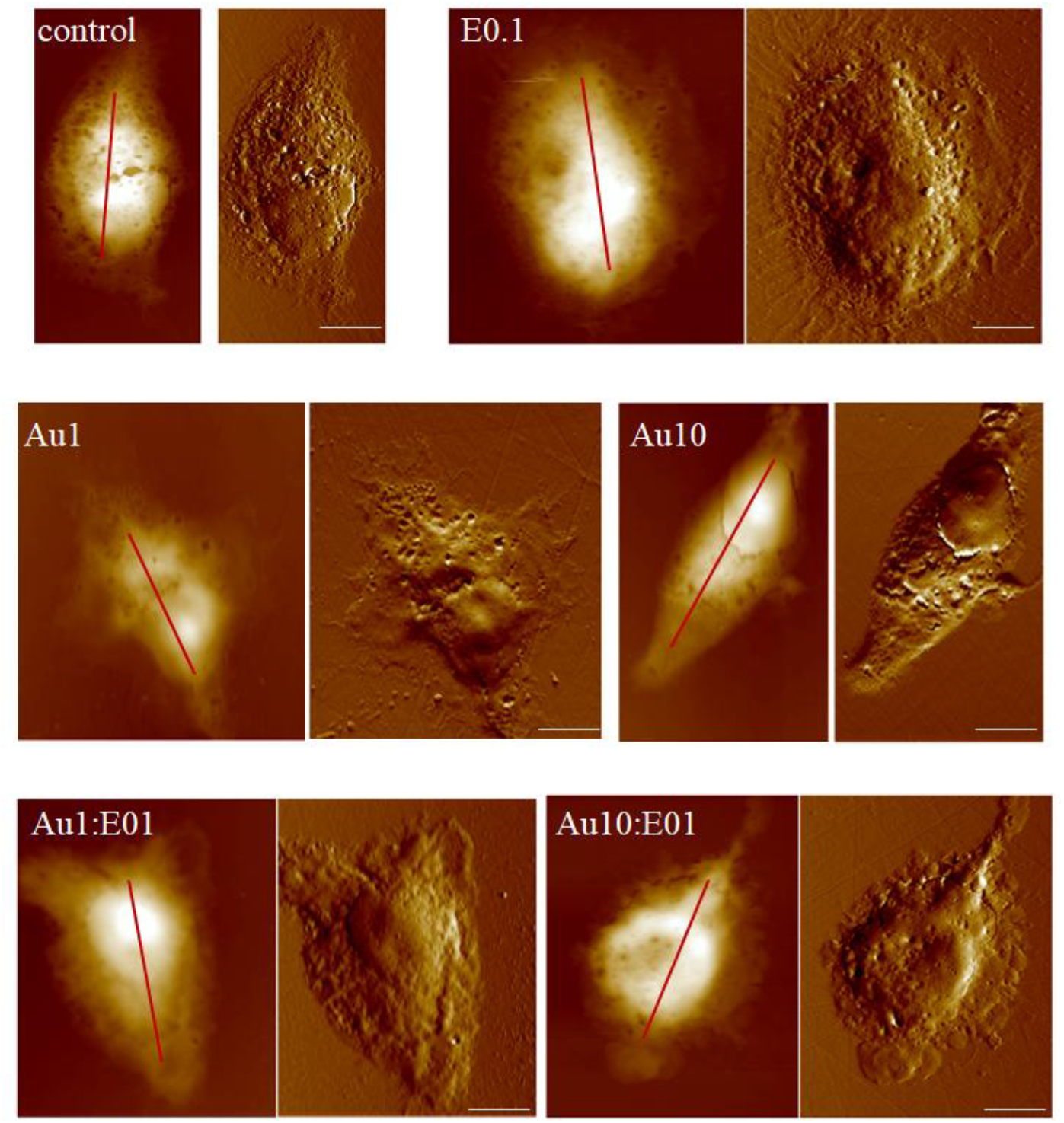
AFM topographies of H1299 cells after 24 h of treatment. Height images (left) and deflection images (right) are shown for untreated cells (control) and cells treated with erlotinib at 0.1 μM (E0.1), AuNPs at 10 mg/L (AuNPs10) and 1 mg/L (AuNPs1), as well as erlotinib–AuNP conjugates containing 10 mg/L AuNPs and 0.1 μM erlotinib (AuNPs10.1) or 1 mg/L AuNPs and 0.1 μM erlotinib (AuNPs1.1). The red straight line indicates the profile along which the measurement points were collected. Scale bar = 10 μm.

Erlotinib:AuNP conjugates are presented in Figure 7. Representative averaged AFM–IR spectra are presented in Figure S2. The biochemical response of the cells following treatment with erlotinib at a concentration of 0.1 µM is shown in Figure 7A. The PC1 loading plot (Fig. 7B) revealed positive bands at 1242 cm^−1^ and 1736 cm^−1^, corresponding to lipid-associated vibrational modes.^22–24^ In contrast, the negative feature observed at 1654 cm^−1^ was assigned primarily to the amide I band of proteins.^25^ This spectral pattern suggests that erlotinib treatment was associated with an increase in cellular lipid content. However, it should be emphasized that, according to the MTS assay results, erlotinib at this concentration (0.1 µM) did not induce a significant cytotoxic effect on cell viability.

**Fig 7.**
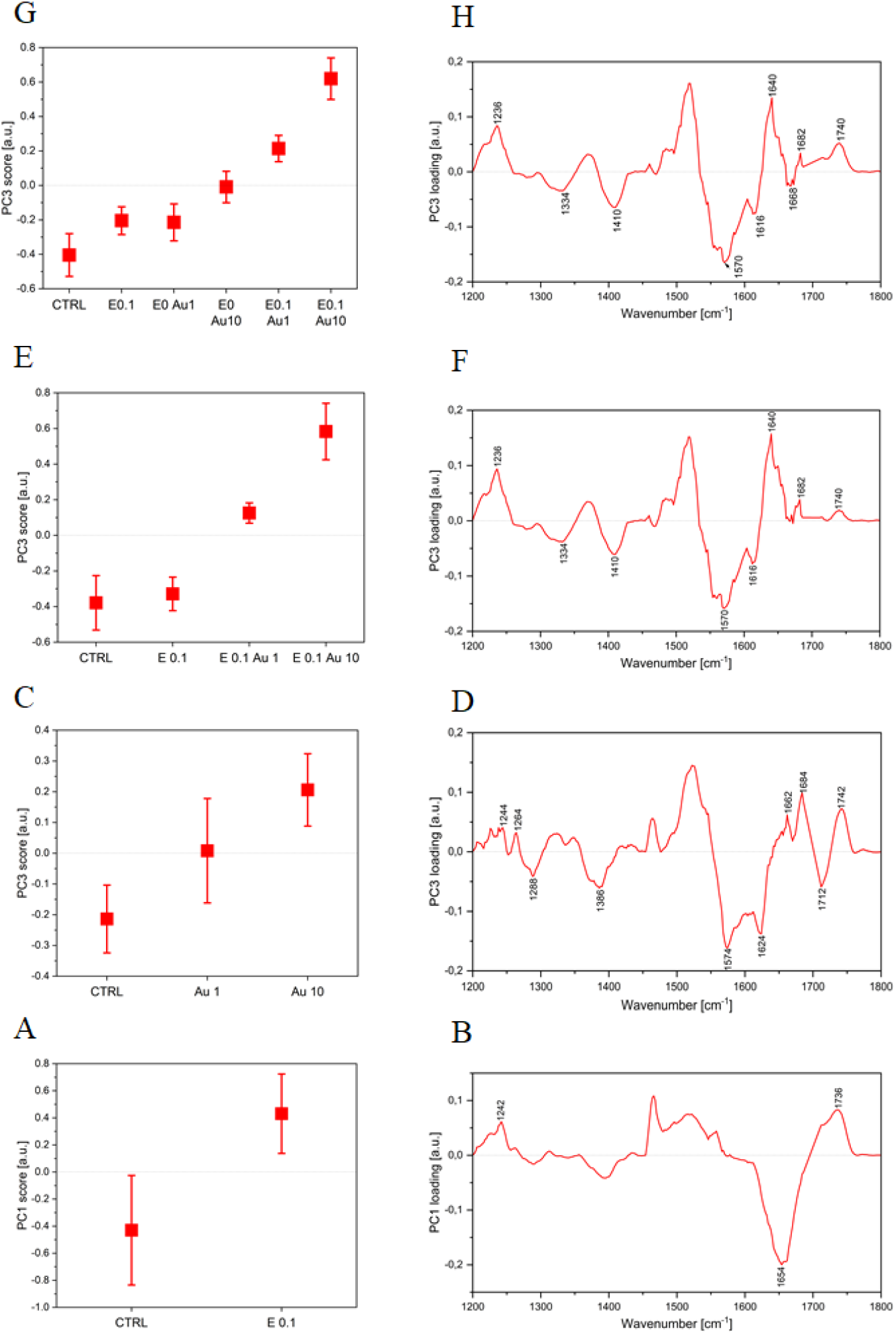
PCA results presented as average score and loading plots derived from AFM–IR spectra of H1299 cells untreated (ctrl) and treated with 0.1 µM erlotinib (E0.1), AuNPs at concentrations of 1 mg/L (A1) and 10 mg/L (A10), and conjugates containing 0.1 µM erlotinib with 1 mg/L AuNPs (E01A1) and 10 mg/L AuNPs with 0.1 µM erlotinib (E01:A10). All samples were incubated for 24 h before measurements.

Considering the effect of AuNPs on the biochemical response of H1299 cells, the PC3 loading plot (Fig. 7D) exhibited a distinct spectral pattern. Positive features observed at 1264 cm^−1^ and 1742 cm^−1^ were attributed to lipids.^22–24^ In addition, bands located at 1662 cm^−1^ and 1684 cm^−1^, assigned to *β*-turn and antiparallel *β*-turn structures within the amide I region, were also detected. Simultaneously, the negative feature at 1624 cm^−1^ was assigned to the *β*-sheet component of the amide I band.^26,27^ These findings suggest that AuNP treatment strongly influenced protein secondary structure, particularly promoting *β*-turn and antiparallel *β*-turn conformations.

Figure 7E illustrates the PC3 loading plot summarizing the results obtained for untreated H1299 cells (control) and cells treated with erlotinib (0.1 µM), as well as with the erlotinib:AuNP systems at concentrations of 0.1 µM:1 mg/L and 0.1 µM:10 mg/L, respectively. The comparison indicates that untreated cells and cells treated with erlotinib alone at 0.1 µM exhibited a similar biochemical profile. Negative bands observed in the loading plot were assigned to the *β*-sheet protein secondary structure at 1616 cm^−1^ (Fig. 7F). In contrast, the spectra corresponding to the investigated erlotinib:AuNP systems displayed positive bands at 1740 cm^−1^, attributed to lipids, as well as at 1640 cm^−1^ and 1682 cm^−1^, corresponding to unordered protein secondary structures and antiparallel *β*-turn conformations, respectively.^26^ These results indicate that the erlotinib:AuNP systems induced noticeable alterations in both lipid composition and protein secondary structure organization in H1299 cells.

Similar conclusions can be drawn from the comparison of results obtained for all investigated conditions, including untreated cells, cells treated with erlotinib (0.1 µM), AuNPs (1 mg/L and 10 mg/L), and the erlotinib:AuNP systems (0.1 µM:1 mg/L and 0.1 µM:10 mg/L). The negative features observed at 1616 cm^−1^, characteristic of untreated cells as well as cells treated with erlotinib and AuNPs alone, were assigned to the *β*-sheet component of protein secondary structure. In contrast, the positive features characterizing the cellular response to the erlotinib:AuNP systems were observed at 1740 cm^−1^, corresponding to lipids, and at 1640 cm^−1^ and 1682 cm^−1^, assigned to unordered protein structures and antiparallel β-turn conformations, respectively. These findings further confirm that the combined erlotinib:AuNP treatment induced pronounced biochemical alterations, particularly affecting lipid content and protein secondary structure in H1299 cells.

## CONCLUCIONS

In this study, erlotinib-functionalized gold nanoparticles (erlotinib:AuNPs) were successfully developed and characterized as a potential nanosystem for targeted intracellular drug delivery. The applied strategy enabled the use of a non-cytotoxic concentration of erlotinib (0.1 µM), which alone did not significantly affect H1299 cell viability, while conjugation with AuNPs resulted in a pronounced reduction of cell viability to approximately 60%. These findings indicate that AuNPs effectively enhanced the intracellular delivery and biological activity of erlotinib.

Fluorescence microscopy confirmed the uptake of the developed nanosystems by H1299 metastatic non-small cell lung cancer cells and demonstrated their preferential localization in the perinuclear and perimitochondrial regions.

The performed 3D RS mapping provided direct spectroscopic confirmation of intracellular nanosystem localization. Owing to the characteristic Raman signatures of erlotinib:AuNPs, cluster analysis enabled precise identification of the nanosystems inside the cells. Moreover, 3D RS mapping exhibited significantly higher sensitivity than fluorescence imaging, allowing detection of intracellular nanosystems even at concentrations for which fluorescence microscopy failed to reveal their presence. The observation of characteristic erlotinib RS bands within the cellular spectra further confirmed successful intracellular transport of the drug-loaded nanosystems. These results demonstrate the high potential of Raman-based imaging for sensitive tracking of nanoparticle-mediated drug delivery systems.

AFM–IR spectroscopy combined with principal component analysis revealed that internalization of the erlotinib/AuNP nanosystems induced pronounced biochemical alterations in H1299 cells. The observed spectral changes indicated modifications in lipid composition together with significant rearrangements in protein secondary structure, particularly involving increased unordered and antiparallel *β*-turn conformations. In contrast, cells treated with erlotinib alone at 0.1 µM displayed a biochemical profile similar to untreated cells, confirming that the detected molecular alterations were predominantly associated with the nanosystem-mediated delivery process.

Overall, the presented results demonstrate that the combination of fluorescence microscopy, 3D RS mapping, and AFM–IR spectroscopy constitutes a powerful multimodal analytical platform for investigating nanoparticle-based drug delivery systems. The developed erlotinib:AuNP conjugates exhibit efficient intracellular delivery, selective uptake by cancer cells, and the ability to induce measurable biochemical changes at the molecular level, highlighting their potential for future theranostic and targeted anticancer applications.

## Conflicts of Interest

The authors declare no conflict of interest.

## Acknowledgments

This work was supported by the National Science Centre Poland (No. 2016/21/D/ST4/02178 to N. P.). The authors would like to thank Ms. Klaudia Cieżak for her assistance with cell culture.

## SUPPORTING INFORMATION

**Fig. S1.**
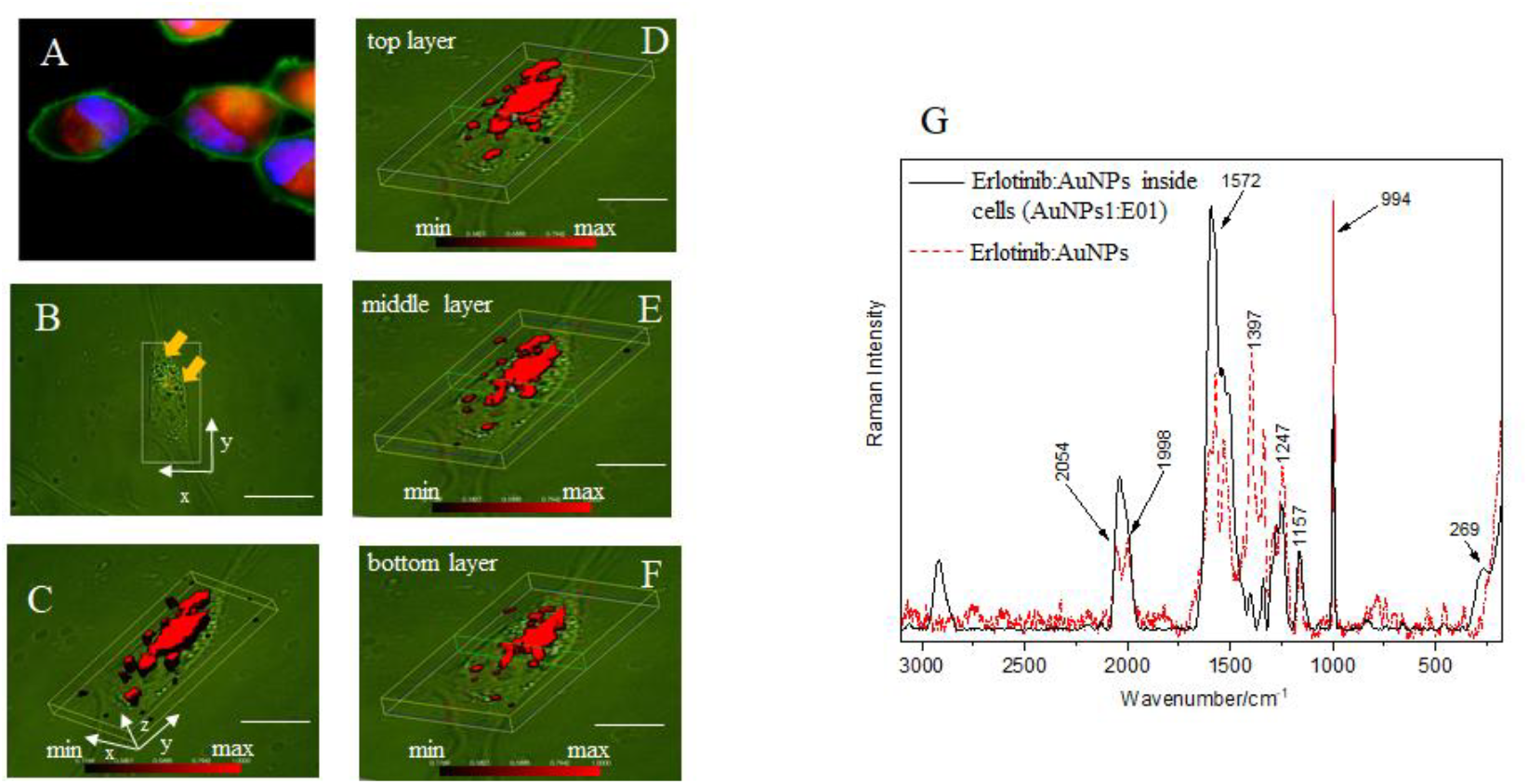
The fluorescent (A) and microscope image (B) of H1299 cells treated for 24 h with AuNP:erlotinib conjugates containing 1 mg/L AuNPs and 0.1 µM erlotinib (AuNPs1:E01). The intracellular distribution of the conjugates, determined by cluster analysis (CA), is shown in panels C–F. Panel G presents the averaged Raman spectrum of the intracellular AuNP:erlotinib cluster (black solid line) compared with the SERS spectrum of the AuNP:erlotinib conjugate recorded after 24 h of adsorption at 37 °C (red dotted line). The arrows indicate the conjugates’ accumulations. Scale bar: 20 µm.

**Fig. S2.**
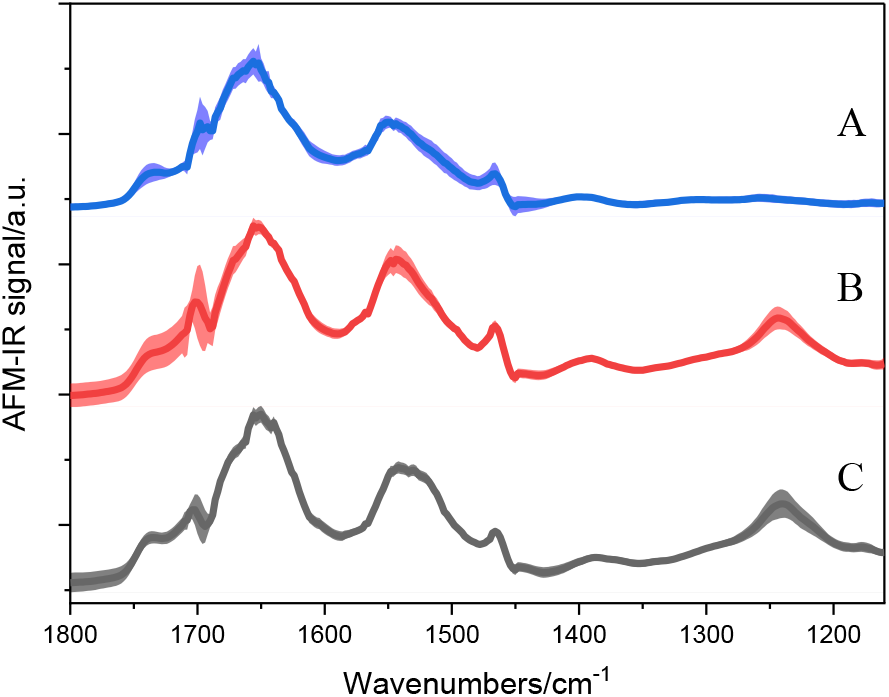
The averaged AFM–IR spectra of spectra of H1299 cells untreated (A) and treated with AuNPs at concentrations of 10 mg/L (B), and conjugates containing 10 mg/L AuNPs with 0.1 µM erlotinib (C). All samples were incubated for 24 h before measurements. The shaded area represents the standard deviation.

## Notes

### Competing Interest Statement

The authors have declared no competing interest.

